# Ultrasound-mediated blood–brain barrier modulation enhances T-cell access but requires immune activation for effective CNS immunity

**DOI:** 10.64898/2026.08.14.744698

**Authors:** Marco Gallus, Akane Yamamichi, Victor Andres Arrieta, Takahide Nejo, Lan Phung, Atsuro Saijo, Pavlina Chuntova, Jianwen Lu, Su Phyu, Heather L. Benway, Aishi Zhao, Kaori Okada, Payal B. Watchmaker, Jeffrey Haegelin, Senthilnath Lakshmanachetty, Karl Habashy, Jacob S. Young, Michael Canney, Roger Stupp, Andres M Salazar, Adam M Sonabend, Hideho Okada

**Affiliations:** UCSF Department of Neurological Surgery, San Francisco, CA, USA; Department of Neurological Surgery, Feinberg School of Medicine, Northwestern University, Chicago, IL, USA; Northwestern Medicine Malnati Brain Tumor Institute of the Lurie Comprehensive Cancer Center, Feinberg School of Medicine, Northwestern University, Chicago, IL, USA; Carthera, Lyon, France; Oncovir, Inc., Washington, D.C, USA; Parker Institute for Cancer Immunotherapy, San Francisco, CA, USA; UCSF Helen Diller Family Comprehensive Cancer Center, San Francisco, CA, USA; Department of Neurology, Feinberg School of Medicine, Northwestern University, Chicago, IL, USA; Division of Hematology/Oncology, Department of Medicine, Feinberg School of Medicine, Northwestern University, Chicago, IL, USA

## Abstract

Immunotherapy shows limited efficacy in brain tumours, where restricted immune access, antigenic heterogeneity and local immunosuppression constrain durable responses. Low-intensity pulsed ultrasound with microbubbles (LIPU+MB) transiently modulates the blood–brain barrier (BBB) and is widely assumed to enhance immunotherapy by facilitating drug and immune cell penetration into the central nervous system (CNS). However, whether increased anatomical access alone is sufficient to generate effective CNS immunity remains unclear. Here, using a transgenic mouse model with astrocyte-restricted antigen expression, we showed that BBB modulation alone is insufficient to generate functional T-cell immunity in the CNS. Although LIPU+MB enabled rapid T-cell entry, accumulation required prior T-cell activation and integrin-dependent mechanisms, indicating that entry remains governed by canonical immune processes. Moreover, T-cells failed to persist owing to insufficient activation of antigen-presenting cells (APCs) within the CNS. Systemic immune adjuvants (poly-ICLC and IL-2; PI) induced APC activation, promoted tissue-resident-memory-like differentiation and supported durable T-cell responses. LIPU+MB further enhanced these responses by increasing T-cell recruitment, resulting in greater accumulation than with PI alone. Mechanistically, antigen presentation by bone marrow–derived APCs was more critical than that by microglia for the accumulation and persistence of antigen-specifc T-cells in the CNS. In antigenically heterogeneous glioma models resistant to CAR T-cell therapy, combining PI with BBB modulation enhanced the efficacy of immunotherapy, which was mirrored by prolonged survival and endogenous tumour-specific T-cell responses, consistent with epitope spreading. Together, these findings define key limitations of LIPU+MB in enabling effective T-cell therapy and establish that BBB modulation must be coupled to systemic immune activation to support T-cell-mediated antitumour immunity in the CNS.

## Main

Adoptive T-cell therapies, including chimeric antigen receptor (CAR) T-cell therapy, have transformed the treatment of select B-cell malignancies (*1–3*), yet their efficacy in malignant brain tumours, most notably glioblastoma (GBM), has remained limited (*4, 5*). Clinical resistance reflects a convergence of challenges, including profound intratumoural heterogeneity, a strongly immunosuppressive tumour microenvironment, and restricted immune access to the CNS (4, 6). While CAR T-cells can effectively eliminate antigen-positive tumour cells, residual antigen-negative subclones frequently persist and drive recurrence, underscoring the need to induce endogenous, polyclonal T-cell responses against a broader repertoire of glioma-associated antigens (7).

Although the CNS is no longer considered strictly immune-privileged (8), immune surveillance of the brain parenchyma remains tightly constrained. Under homeostatic conditions, T-cell trafficking is largely restricted to meningeal and choroid plexus compartments, while the BBB limits immune entry into the parenchyma (8–11). In GBM, BBB disruption is spatially heterogeneous: contrast-enhancing tumour regions often exhibit permeability, whereas infiltrative margins and many diffuse midline and lower-grade gliomas retain an intact BBB (11, 12). These regions are precisely where residual tumour persists and where immunotherapies frequently fail (4, 13, 14).

Consistent with this, vaccine-induced tumour-reactive T-cells can be detected systemically, but often fail to accumulate within the tumour microenvironment (15–17).

Low-intensity pulsed ultrasound with microbubbles (LIPU+MB) enables transient, spatially controlled modulation of BBB permeability(18, 19) and has emerged as a promising strategy to enhance immunotherapy delivery to the CNS (20–24). However, whether LIPU+MB-mediated BBB modulation alone is sufficient to generate functional antigen-specific T-cell immunity in the CNS remains unclear, and the mechanisms limiting the efficacy of ultrasound-enabled immunotherapy are poorly defined (25).

We hypothesized that the efficacy of LIPU+MB in enabling T-cell responses maybe constrained by the need to coordinate two mechanistically distinct processes: spatial immune access and immunological licensing. Using a transgenic mouse model with astrocyte-restricted antigen expression and an intact BBB, we show that ultrasound-mediated BBB modulation permits transient T-cell entry but is insufficient to sustain immunity owing to inadequate antigen-presenting cell activation. However, when combined with systemic immune activation, BBB modulation enhances T-cell recruitment and furthermore promotes durable antitumour responses in a heterogeneous glioma model resistant to CAR T-cell therapy. Together, our findings define key mechanistic limitations of LIPU+MB in enabling effective T-cell therapy in the CNS and establish a framework for integrating BBB modulation with immune licensing to improve immunotherapy in brain tumours.

## Results

### LIPU+MB permits transient antigen-specific T-cell entry, but remains dependent on canonical trafficking mechanisms

Recent studies, including our own (22, 23), have suggested that LIPU+MB enhances CAR T-cell trafficking into the central nervous system CNS (20, 21). However, the underlying mechanism and whether ultrasound-mediated BBB modulation alone is sufficient to generate durable antigen-specific immunity within the CNS remained unclear.

To address this question, we generated a novel transgenic mouse model in which defined MHC class I and class II-restricted viral and tumour epitopes (gp100, LCMVgp33-41, LCMVgp61-80, TRP-1) are selectively expressed by astrocytes under control of the GFAP promoter (GFAP-minigene mice, **Fig. 1a**).

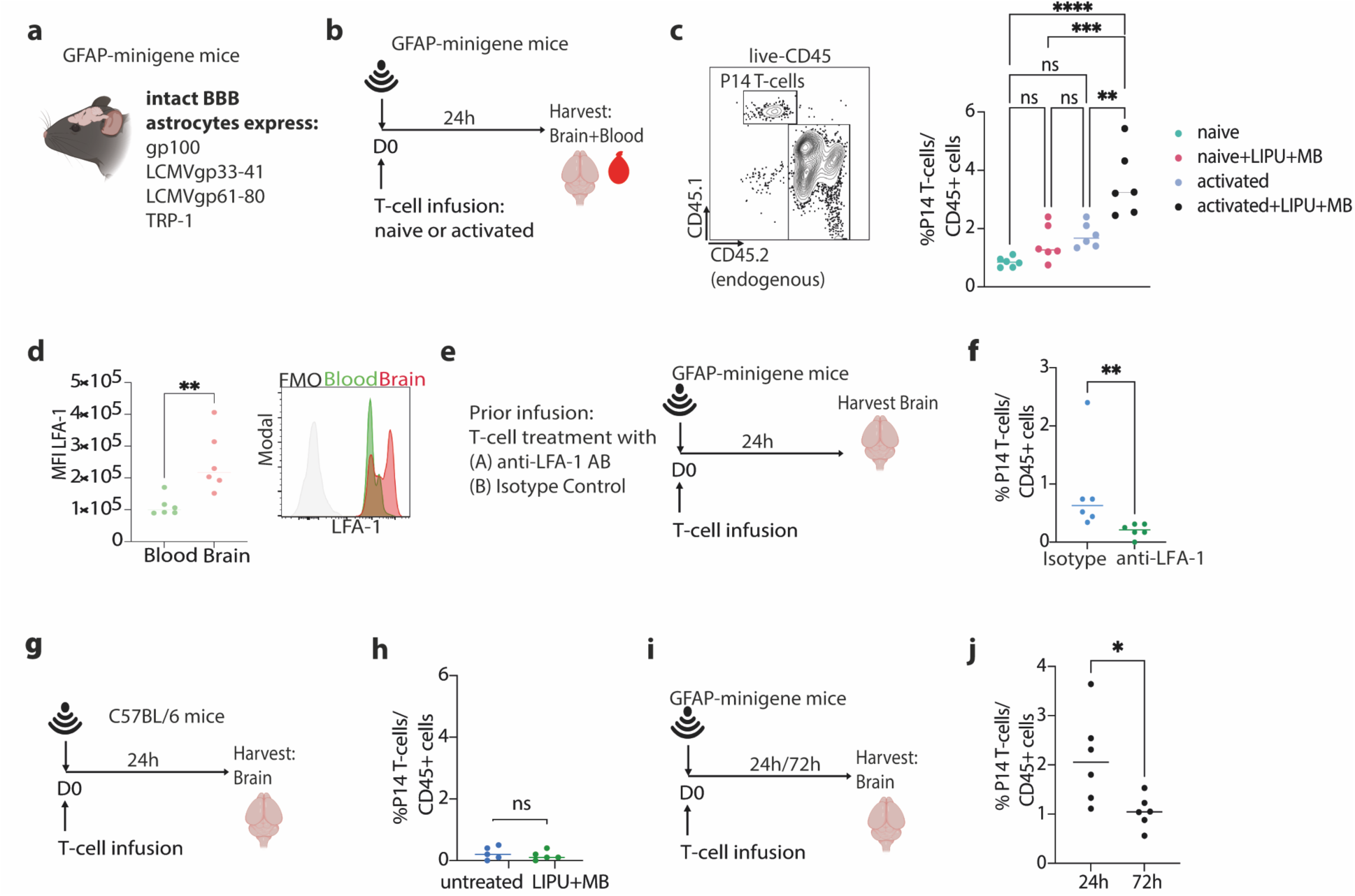
LIPU+MB enables transient antigen-specific T-cell entry but not durable CNS immunity. **a**, GFAP–minigene mice expressing astrocyte-restricted model antigens (gp100, LCMVgp33–41, LCMVgp61–80 and TRP-1) under intact BBB conditions. **b**, Experimental design. Naïve or ex vivo activated P14 CD8⁺ T-cells were adoptively transferred into GFAP–minigene mice followed by LIPU+MB treatment. Brains were analysed 24 h later. **c**, Representative gating strategy and quantification of transferred P14 T-cells among CD45⁺ brain leukocytes in mice receiving naïve or activated T-cells with or without LIPU+MB treatment. **d**, Representative histograms and quantification of LFA-1 expression on transferred P14 T-cells recovered from blood and brain 24 h after transfer. **e**, Experimental design for LFA-1 blockade. Activated P14 T-cells were pre-incubated with anti–LFA-1 blocking antibody or isotype control before adoptive transfer and LIPU+MB treatment. **f**, Quantification of transferred P14 T-cells among CD45⁺ brain leukocytes following LFA-1 blockade or isotype control treatment. **g**, Experimental design assessing CNS trafficking in wild-type C57BL/6 mice lacking cognate antigen expression. **h**, Quantification of transferred P14 T-cells among CD45⁺ brain leukocytes in untreated or LIPU+MB-treated wild-type mice. **i**, Experimental design assessing persistence of transferred T-cells following LIPU+MB treatment in GFAP–minigene mice. Brains were analysed 24 or 72 h after transfer. **j**, Quantification of transferred P14 T-cells among CD45⁺ brain leukocytes at 24 and 72 h after transfer. Each symbol represents one biologically independent mouse; n =5-6 biologically independent mice per group. Statistical significance was determined using one-way ANOVA with Tukey’s multiple-comparisons test or two-tailed unpaired t-tests. *P < 0.05, **P < 0.01, ***P < 0.001, ****P < 0.0001; ns, not significant.

Commonly used orthotopic glioma models exhibit early and heterogeneous BBB disruption (**Extended Data Fig. 1**), thereby preventing us from studying precise immune mechanisms under intact BBB conditions. On the other hand, the GFAP-minigene mice preserve physiological barrier integrity and model clinically relevant settings in which the BBB remains intact.

To determine whether ultrasound-enabled CNS recruitment depends on physiological trafficking mechanisms, activated or naïve LCMV gp33-specific P14 CD8⁺ T-cells were adoptively transferred into GFAP–minigene mice followed by LIPU+MB treatment **(Fig. 1b)**. Under physiological and inflammatory conditions, leukocyte trafficking into the CNS is regulated through a multistep adhesion cascade involving integrin-mediated interactions with the cerebrovascular endothelium. In particular, the integrin LFA-1 (CD11a/CD18), expressed on activated T-cells, binds endothelial ICAM-1 to promote adhesion and diapedesis (10, 26, 27). Consistent with this framework, adoptive transfer of activated T-cells resulted in significantly greater CNS accumulation than transfer of naïve (CD44⁻) T-cells following LIPU+MB **(Fig. 1c)**. Moreover, adoptively transferred T-cells recovered from the CNS exhibited significantly higher LFA-1 expression compared to those in the periphery **(Fig. 1d)**. To determine whether this enhanced accumulation reflected activation-associated trafficking rather than passive entry through a mechanically modulated BBB, we next blocked LFA-1 on T-cells before intravenous infusion. LFA-1 blockade markedly reduced CNS accumulation following LIPU+MB treatment **(Fig. 1e,f)**, indicating that ultrasound-enabled entry remains dependent on canonical trafficking mechanisms.

Importantly, adoptive transfer of activated antigen-specific T-cells into wild-type C57BL/6 mice resulted in only minimal CNS accumulation following LIPU+MB **(Fig. 1g,h)**, indicating that antigen availability further contributes to effective T-cell recruitment and retention within the CNS. Furthermore, although adoptive transfer of activated P14 CD8⁺ T-cells followed by LIPU+MB resulted in rapid CNS accumulation at 24 h, T-cell frequencies returned to near-baseline levels by 72 h, indicating a failure to sustain antigen-specific immunity within the CNS **(Fig. 1i,j)**. Together, these findings demonstrate that LIPU+MB-mediated BBB modulation permits transient antigen-specific T-cell entry but does not bypass physiological trafficking requirements or sustain durable CNS immunity.

### Durable CNS T-cell immunity requires immune licensing beyond LIPU+MB-enabled access

Having established that LIPU+MB-mediated T-cell entry is activation-dependent but fails to result in sustained CNS accumulation, we next asked whether the use of adjuvants, such as those activating the host immune system, including antigen-presenting cells, is required to license durable T-cell responses following ultrasound-mediated BBB opening.

To this end, non-activated P14 T-cells were adoptively transferred into GFAP-minigene mice followed by repeated LIPU+MB treatments, with or without concurrent administration of poly-ICLC and IL-2 (PI) (**Fig. 2a**) as immune adjuvants. Poly-ICLC, a synthetic double-stranded RNA and potent TLR3/MDA5 agonist, primarily activates innate immune pathways, leading to type I interferon induction, proinflammatory cytokine release, and maturation of dendritic cells (DCs) (*15, 28, 29*). IL-2 acts directly on T-cells, promoting their proliferation, differentiation, and survival while augmenting effector functions, such as granzyme-and perforin-mediated cytotoxicity (*30*).

**Fig 2.**
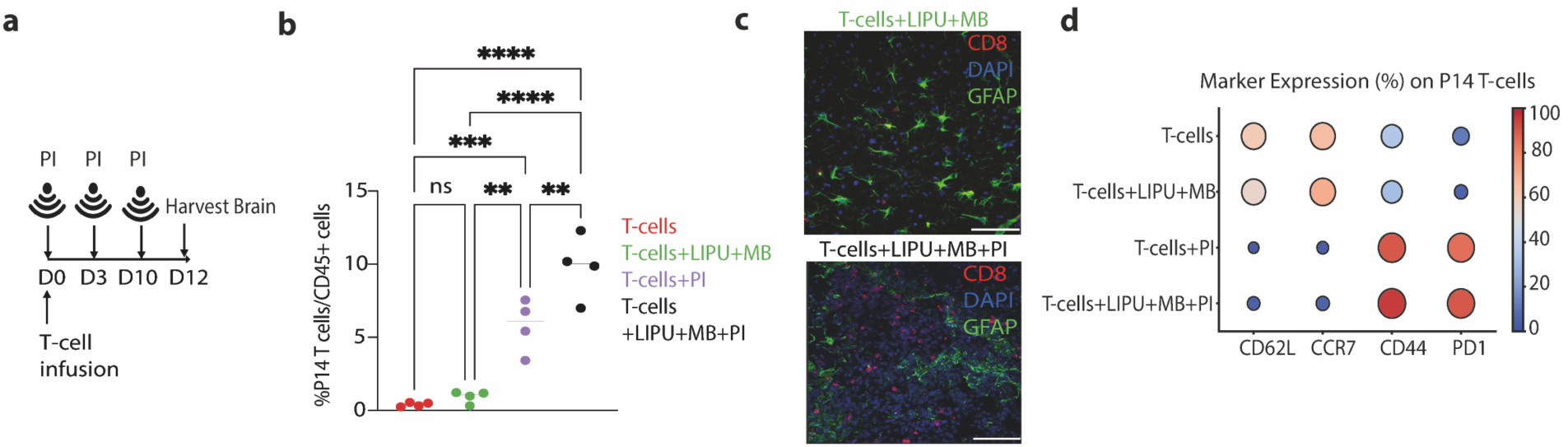
PI promotes the persistence of antigen-specific T-cells in the CNS. **a,** Treatment schedule including repeated LIPU+MB and/or PI treatment. Brains were analyzed at D12. **b,** Frequency of transferred P14 T-cells among CD45⁺ brain leukocytes at D12. Each dot represents one mouse; n = 4 per group. Representative of two independent experiments **c,** Representative immunofluorescence images of brain sections of LIPU+MB or LIPU+MB+PI treated animals (CD8, red; GFAP, green; DAPI, blue). **d,** Bubble plot indicates percentage of marker expression on adoptively transferred cells identified in the brain; n = 4 mice per group. Statistical significance was determined using one-way ANOVA with Tukey’s multiple comparisons test (****P < 0.0001; ***P < 0.001; **P < 0.01; ns, not significant).

LIPU+MB alone, even when repeated three times, did not induce sustained T-cell accumulation, with transferred cells remaining at levels comparable to untreated controls **(Fig. 2b,c)**. In contrast, PI administration promoted robust accumulation of adoptively transferred antigen-specific T-cells within the brain (**Fig. 2b,c)**. A combination of PI with additional LIPU+MB treatment resulted in greater T-cell accumulation than PI alone, indicating additive effects.

Furthermore, PI treatment promoted differentiation of CNS-infiltrating T-cells toward a CD44⁺CD62L⁻CCR7^-^CD69⁺ tissue-resident memory–like phenotype **(Fig. 2d).**

Together, these findings demonstrate that ultrasound-mediated BBB modulation can potentiate CNS-antigen-specific T-cell recruitment but depends on host immune activation.

### LIPU+MB recruits myeloid cells but does not license antigen-presenting cells

PI treatment promotes durable immunity, whereas LIPU+MB alone fails to sustain long-term T-cell persistence. To elucidate the mechanisms underlying this limitation, we next examined how LIPU+MB influences antigen presentation within the CNS. Given that sustained T-cell responses depend on effective antigen presentation and costimulatory signaling in a pro-inflammatory environment, we assessed the impact of LIPU+MB on CNS-resident antigen-presenting cells and their capacity to support T-cell activation.

Both LIPU+MB treatment and PI treatment increased the frequency of CD45^high^-CD11b^+^ myeloid cells in the CNS compared to untreated controls, consistent with enhanced recruitment across the BBB (**Extended Data Fig. 2**). However, we observed no statistically significant differences comparing LIPU+MB-treated animals with those receiving LIPU+MB in combination with PI (**Extended Data Fig. 2**).

We hypothesized that LIPU+MB-mediated BBB opening provides anatomical access for CD45^high^-CD11b^+^ myeloid cells but fails to ensure their maturation to adequate antigen-presenting cells within the intrinsically tolerogenic CNS microenvironment. To identify candidate pathways underlying this distinction, we performed bulk RNA sequencing of brains derived from GFAP-minigene mice 24 hours after treatment with LIPU+MB alone or in combination with PI (**Extended Data Fig. 3a,b)**. Relative to LIPU+MB alone, combination therapy induced a robust antigen-presenting transcriptional program characterized by upregulation of genes encoding components of the antigen-processing and presentation machinery (such as *H2-K1, H2-D1, B2m, Cd86, Cd274, Icosl, Tap1, Tap2, Tapbp, Psmb8 and Psmb9)* and costimulatory molecules *(such as Cd86, Cd274, Icosl)* (**Extended Data Fig. 3c)**. These data suggest that the immune adjuvant treatment induced a more robust antigen-presenting programme than LIPU+MB alone and motivated subsequent phenotypic validation at the protein level by flow cytometry (**Fig. 3a-c**). Spectral flow cytometry of CNS-resident microglia and infiltrating myeloid cells indicated that, relative to untreated controls, LIPU+MB alone modestly increased CD80, CD86, and MHC class II expression on microglia and was associated with a proportional decrease in these markers on CD11b^+^-CD45^high^ myeloid cells. In contrast, immune adjuvant treatment, whether administered alone or in combination with LIPU+MB; markedly upregulated CD80, CD86, MHC class I, and MHC class II expression on CNS-resident microglia and, to a lesser extent, on myeloid populations **(Fig. 3c)**.

**Fig. 3.**
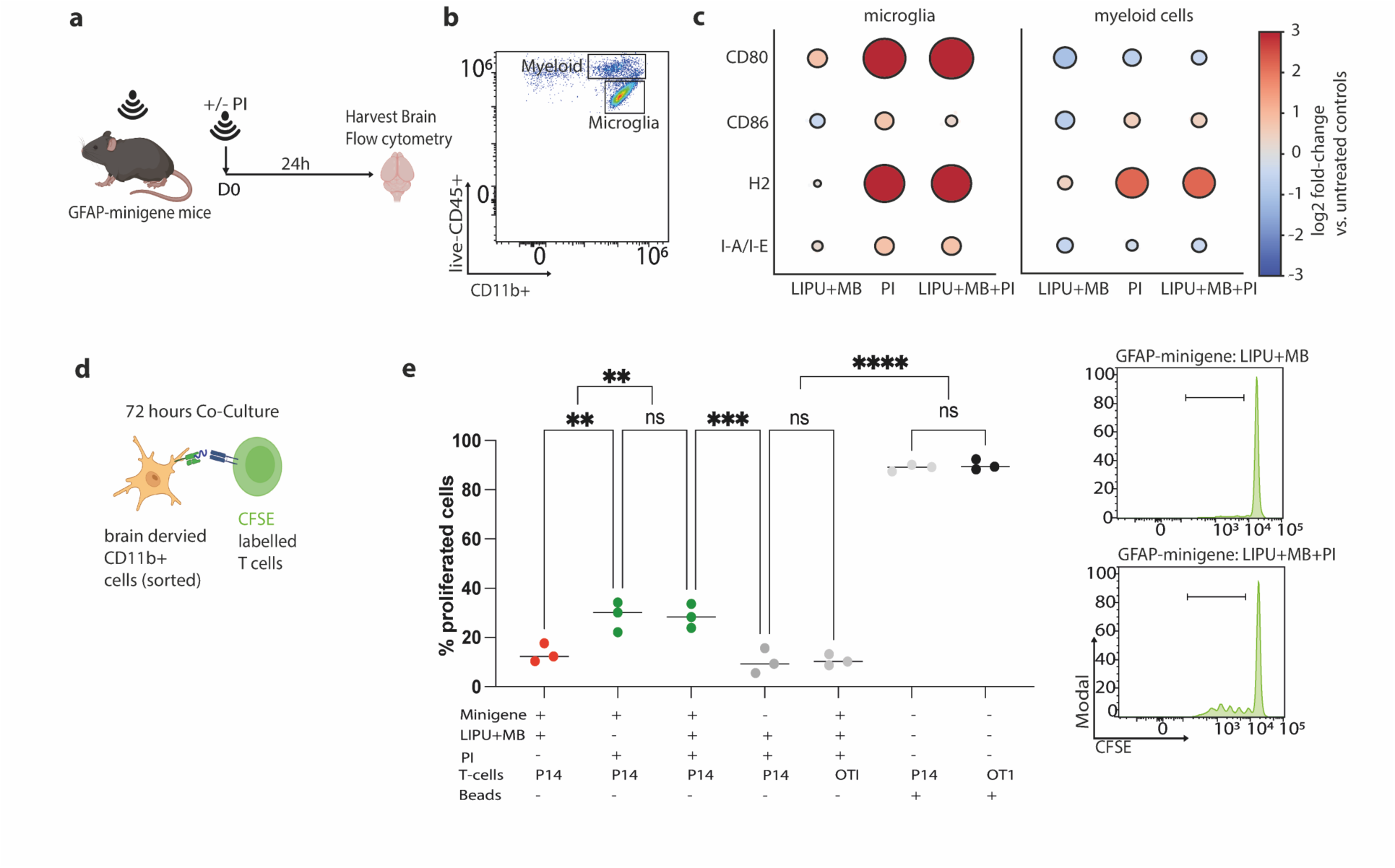
Immune adjuvants, but not LIPU+MB, induce antigen-presenting programs in CNS myeloid cells. **a,** Experimental design. GFAP–minigene mice were treated with LIPU+MB with or without PI. Brains were harvested and analysed by flow cytometry. **b,** Representative gating strategy identifying brain-resident microglia (CD11b⁺CD45^int^) and infiltrating myeloid cells (CD11b⁺CD45^high^) within CD45⁺ brain leukocytes. **c,** Summary of antigen presentation marker expression in microglia and infiltrating myeloid cells following treatment. Circle size and colour intensity denote log₂ fold change relative to untreated controls. **d,** Experimental design for ex vivo T-cell priming assay. Brain-derived CD11b⁺ myeloid cells were isolated and co-cultured with CFSE-labelled antigen-specific P14 T-cells or control OT-I T-cells. **e,** Quantification of T-cell proliferation under the indicated conditions; n = 3 per group. Representative of two independent experiments. Representative CFSE dilution histograms are shown. Statistical significance was determined using one-way ANOVA with multiple comparisons (****P < 0.0001; ***P < 0.001; **P < 0.01; ns, not significant).

Functionally, CD11b⁺ cells isolated from GFAP–minigene brains after combined LIPU+MB and PI treatment induced significantly greater proliferation of antigen-specific P14 T-cells compared with LIPU+MB alone using an ex-vivo co-culture assay. No proliferation was observed in antigen-mismatched OT-I T-cells, confirming antigen specificity (**Fig. 3d-e**).

Together, these findings demonstrate that LIPU+MB enhances myeloid cell recruitment but fails to induce a pro-inflammatory, antigen-presenting phenotype, thereby limiting effective T-cell activation within the CNS.

### Antigen-presentation by bone-marrow-derived cells determines CNS T-cell accumulation following LIPU+MB+PI treatment

Bone marrow–derived cells have been proposed as key gatekeepers for CNS T-cell trafficking (9, 31–36). To test the role of antigen presentation, especially by bone marrow–derived myeloid cells, in driving effective T-cell immunity after LIPU+MB+PI, we generated bone marrow chimeric (BMC) mice in which major histocompatibility complex class I (MHC-I) expression was selectively ablated in hematopoietic cells but preserved in radioresistant CNS-resident microglia. GFAP–minigene mice were lethally irradiated and reconstituted with β2-microglobulin–deficient bone marrow (B2M-BMC), resulting in loss of MHC-I expression selectively on bone marrow–derived cells **(Fig. 4a, Extended Data Figure 4).**

**Fig. 4.**
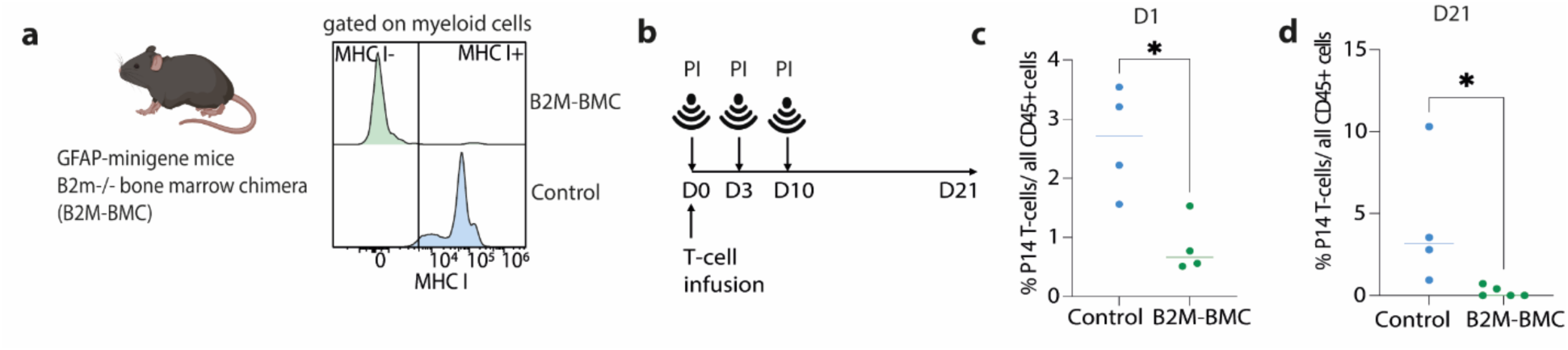
Bone marrow–derived MHC class I expression is required for LIPU+MB+PI-mediated T-cell accumulation in the brain. **a,** Experimental design. Bone marrow chimeric mice were generated by transplantation of β2-microglobulin–deficient (B2m⁻/⁻) bone marrow into irradiated GFAP–minigene recipients (B2M-BMC). **b,** Treatment schedule. Following adoptive transfer of antigen-specific P14 CD8⁺ T-cells at D0, mice received PI on days 3 and 10 and underwent LIPU+MB treatment. Brains were analyzed at day 1 and day 21. **c,** Frequency of transferred P14 T-cells among CD45⁺ brain leukocytes at day 1 after treatment in control B2M-WT-BMC and B2M KO-BMC mice. Each dot represents one mouse; n = 4 per group. **d,** Frequency of transferred P14 T-cells among CD45⁺ brain leukocytes at day 21. Each dot represents one mouse; n = 4/5 per group. Statistical significance was determined using two-tailed t-tests (*P < 0.05).

Following immune reconstitution, mice received adoptive transfer of antigen-specific P14 T-cells combined with LIPU+MB and PI (**Fig. 4b**). Early analysis revealed a significant reduction in the frequency of transferred P14 T-cells in the brains of B2M-BMC compared with control animals (chimeras that received β2-microglobulin–competent bone marrow) at day 1 after treatment (**Fig. 4c**). This defect persisted, as B2M-BMC mice also exhibited markedly reduced accumulation of antigen-specific P14 T-cells in the brain at day 21 (**Fig. 4d**).

These findings demonstrate that antigen presentation by bone marrow–derived cells is required for the persistent accumulation of antigen-specific T-cells in the CNS. Despite local antigen expression and activation of CNS-resident microglia, hematopoietic antigen-presenting cells are therefore essential for establishing durable antigen-specific immunity in the brain. These results are consistent with a model in which bone marrow–derived antigen-presenting cells, including those positioned at CNS interfaces, function as critical gatekeepers of T-cell activation and entry(9, 31–33).

### Combined immune licensing and BBB modulation promote endogenous tumour-specific T-cell responses and prolong survival in antigenically heterogeneous glioma

We next tested whether the combination treatment of LIPU+MB with PI can overcome therapeutic resistance in antigenically heterogeneous tumours. We established an intracranial SB28 glioma model comprising mixed EGFRvIII-positive (80%) and EGFRvIII-negative (20%) tumour cells **(Fig. 5a)**, recapitulating clinically relevant antigenic heterogeneity. SB28 tumours are known to be particularly immunosuppressive and resistant to checkpoint inhibition therapy, resembling human malignant gliomas (37). To enable tracking of endogenous tumour-specific responses, both EGFRvIII-positive and EGFRvIII-negative SB28 cell lines were engineered to express the model antigens gp100 and LCMV gp61–80 **(Fig. 5b).** In tumours uniformly expressing EGFRvIII, a single infusion of EGFRvIII-specific CAR T-cells induced durable tumour control in a substantial fraction of animals **(Extended Data Fig. 5)**. By contrast, in heterogeneous tumours, CAR T-cell therapy alone conferred only limited survival benefit, with consistent relapse **(Extended Data Fig. 5)**, suggesting antigen escape as the dominant resistance mechanism.

**Fig. 5.**
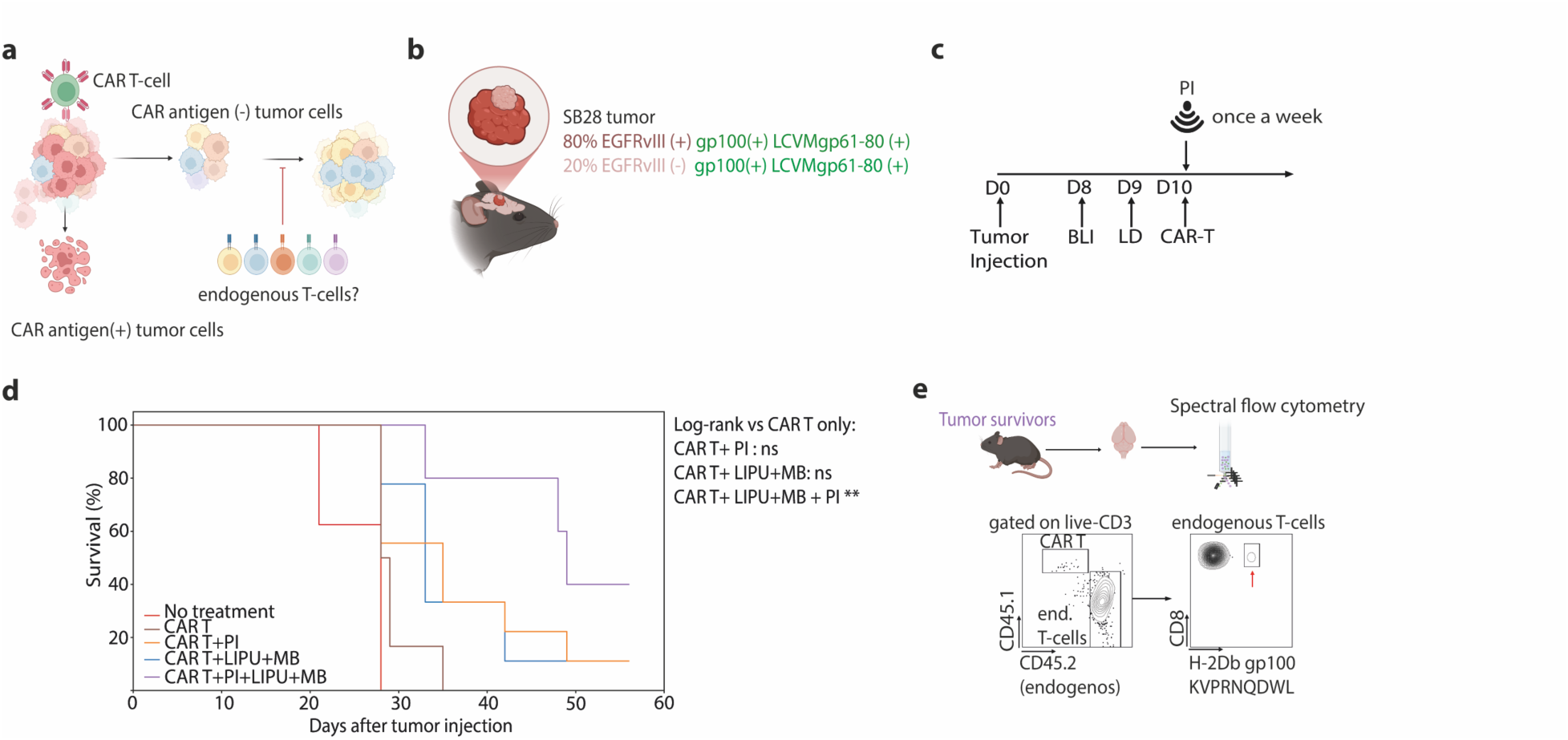
Combined immune licensing and BBB modulation promote endogenous antitumour immunity in antigenically heterogeneous glioma. **a,** Conceptual schematic. CAR T-cells eliminate antigen-positive tumour cells, whereas systemic immune activation promotes recruitment and priming of endogenous T-cells, enabling broader antitumour immunity within the CNS. **b,** Experimental model. Intracranial SB28 gliomas with mixed antigen expression (EGFRvIII⁺ and EGFRvIII⁻ subclones) to model antigen escape. **c,** Treatment schedule. Mice received intracranial tumour implantation, followed by intravenous CAR T-cell transfer, LIPU+MB, and PI. **d,** Kaplan–Meier survival analysis of tumour-bearing mice across treatment groups. Combined therapy resulted in the most durable survival benefit. Significance was assessed by log-rank Mantel–Cox test, **P < 0.01; ns, not significant). **e,** Immune profiling of long-term survivors. Spectral flow cytometry of brain-infiltrating leukocytes demonstrating endogenous tumour-specific T-cell responses (for example, gp100-specific tetramer-positive CD8⁺ T-cells).

We next evaluated whether PI or ultrasound-mediated BBB opening (LIPU+MB) could improve therapeutic efficacy. Both CAR T + PI and CAR T + LIPU+MB resulted in modest survival extensions; however, the majority of the animals ultimately relapsed. In contrast, the combination of CAR T-cells, PI, and LIPU+MB produced the most pronounced survival benefit and generated long-term survivors, consistently outperforming all dual-modality regimens, without showing any clinical evidence of off-tumour toxicity **(Fig. 5c-d)**.

Flow cytometric analysis of the brains of long-term survivors revealed induction of endogenous T-cell responses against gp100, a shared antigen expressed across tumour subclones, consistent with treatment-induced epitope spreading beyond the CAR target **(Fig. 5e)**. In line with this, the therapeutic benefit of combined BBB modulation and immune activation depended on the presence of the gp100 epitope, with significantly enhanced efficacy observed in antigen-positive compared to gp100-negative tumours (**Extended Data Fig. 6).** Together, these findings demonstrate that combining BBB modulation with immune licensing not only enhances therapeutic efficacy but also promotes endogenous, antigen-diversified T-cell responses capable of overcoming tumour heterogeneity.

## Discussion

In this study, we refine the role of LIPU+MB-mediated BBB modulation in the context of cellular immunotherapy in the CNS. Using complementary transgenic and tumour models, we show that LIPU+MB facilitates T-cell access to the CNS but, on its own, does not support durable antigen-specific immunity. Instead, its effects depend on additional coordinated processes, including T-cell activation state and antigen-presenting-cell function. In this setting, LIPU+MB enhances T-cell recruitment when effective immune activation is present. Together, these findings place BBB modulation within the context of established immune mechanisms and help clarify how it can be used to improve the efficacy of cellular immunotherapies.

LIPU+MB enables transient and spatially controlled modulation of BBB permeability and has been shown to enhance the delivery of drugs and immune cells to the brain in preclinical and early clinical studies (20, 22–24, 38). Previous work, including our own, has further demonstrated the induction of a mild, sterile inflammatory response, with upregulation of pro-inflammatory pathways within hours after LIPU+MB (23, 39). However, whether this is sufficient to initiate or sustain antigen-specific adaptive immunity within the CNS has remained unclear.

Using the GFAP–minigene model, which preserves BBB integrity and minimizes tumour-associated confounders, we specifically evaluated the contribution of ultrasound-mediated spatial access. In this setting, LIPU+MB enabled rapid T-cell entry into the CNS but did not support the establishment of a functionally competent antigen-presenting environment. Although myeloid cell recruitment increased, LIPU+MB alone did not induce robust upregulation of antigen presentation or co-stimulatory pathways in microglia or infiltrating myeloid cells, and T-cells failed to persist. Although prior studies have observed that focused ultrasound–mediated BBB opening can transiently activate brain-resident myeloid cells and enhance antitumour immune responses when combined with chemotherapy or checkpoint blockade (20, 22, 24), these studies did not distinguish whether improved immune responses resulted from enhanced anatomical access, ultrasound-induced inflammation, or secondary systemic immune activation. By directly separating these components, our findings demonstrate that LIPU+MB-mediated BBB modulation alone is insufficient to establish a functional antigen-presenting niche and requires an independent immune licensing step. Together, these results indicate that LIPU+MB provides anatomical access but does not, in itself, induce the immune activation required for effective CNS immunity.

Our data indicate that the efficacy of LIPU+MB depends on pre-existing immune activation, involving both APCs and endogenous or adoptively transferred T-cells. CNS recruitment following LIPU+MB depended on T-cell activation status and integrin-dependent mechanisms. Activated T-cells accumulated more efficiently than naïve cells, and LFA-1 blockade reduced CNS accumulation following LIPU+MB. Together with prior observations that ultrasound upregulates endothelial adhesion molecules such as ICAM-1 (20), these findings support a model in which LIPU+MB renders the cerebrovascular interface permissive but does not substitute for activation-dependent trafficking processes. Consistent with this, LIPU+MB enhanced T-cell accumulation in the context of systemic immune activation, resulting in greater CNS infiltration compared with immune activation alone. These findings establish that LIPU+MB amplifies ongoing immune responses rather than initiating adaptive immunity, and that effective T-cell recruitment to the CNS depends on prior T-cell activation and immune licensing.

Using bone marrow chimeric mice, we observed reduced T-cell accumulation when MHC class I expression was absent in hematopoietic cells, despite preserved antigen expression in CNS-resident microglia.

These findings align with emerging evidence that effective T-cell priming in CNS-directed immune responses often depends on antigen presentation outside the brain parenchyma, including by bone marrow–derived antigen-presenting cells at CNS interfaces or in draining lymphoid tissues (9, 34, 40, 41). In this context, our data are consistent with a role for hematopoietic APCs in generating trafficking-competent T-cells capable of responding to ultrasound-enabled access.

At the same time, these results do not exclude contributions from CNS-resident antigen-presenting cells, but suggest that peripheral or interface-associated bone marrow-derived APCs represent an important component of the system required for sustained antigen-specific immunity in the CNS.

Antigen escape has been identified as a dominant mechanism of resistance to CAR T-cell therapy in glioblastoma (7, 14). In our antigenically heterogeneous tumour model, CAR T-cells achieved durable control in antigen-uniform tumours but failed in mixed populations, recapitulating clinical relapse patterns. Combination of LIPU+MB with PI produced a pronounced survival benefit and generated long-term survivors.

Therapeutic efficacy was associated with endogenous T-cell responses against non–CAR-targeted tumour antigens and evidence of epitope spreading. Efficacy was higher in antigen-positive tumour settings, supporting a model in which LIPU+MB+PI promotes recruitment of functionally competent, antigen-specific T-cells. This diversification of the T-cell repertoire provides a mechanistic basis for durable tumour control in heterogeneous disease and aligns with prior observations linking epitope spreading to improved outcomes in cancer immunotherapy (42, 43). These findings indicate that LIPU+MB-mediated spatial immune access alone is insufficient for effective CNS immunotherapy. Clinical studies have demonstrated that focused ultrasound can safely and transiently modulate BBB permeability in patients (24, 38, 44, 45). Our data now suggest that barrier opening must be coupled with systemic immune activation to achieve durable biological and therapeutic effects. In this framework, LIPU+MB is most likely to be effective when integrated with therapies that generate or expand tumour-reactive lymphocytes.

Several limitations warrant consideration. Although the GFAP–minigene model enabled precise mechanistic dissection under conditions of intact BBB integrity, it does not fully recapitulate the cellular and spatial complexity of glioblastoma. The SB28 model captures antigenic heterogeneity and therapeutic resistance; however, engineering tumour cells to express model antigens increases immunogenicity relative to native disease.

Combination treatment with LIPU+MB, PI and CAR T-cells was not associated with overt clinical evidence of off-target toxicity. Nevertheless, future translational development will require careful optimization of dose, timing, and sequencing to define a therapeutic window that maximizes efficacy while limiting excessive inflammation.

Finally, although our data identify activation-dependent and integrin-dependent trafficking as key determinants of ultrasound-enabled immune deployment, additional vascular and stromal factors are likely to contribute and remain to be defined.

Together, our findings redefine the role of LIPU+MB in immunotherapy. While ultrasound-mediated BBB modulation resolves anatomical exclusion, it is insufficient to drive antigen-specific T-cell immunity because it does not induce immune licensing. Instead, LIPU+MB functions as a spatial amplifier of pre-existing immune responses, enhancing recruitment of activated T-cells without initiating adaptive immunity. Consequently, its therapeutic efficacy depends on integration with immune-activating strategies, supporting recruitment and diversification of CNS-tumour-reactive T-cells, thereby enabling durable antitumour immunity in antigenically heterogeneous glioma.

## Materials and Methods

### Study approval

All procedures complied with an Institutional Animal Care and Use Committee–approved protocol. Our study examined male and female animals, and similar findings are reported for both sexes.

### Mice

Mice (8–10 weeks old) were used for all experiments. C57BL/6N (JAX 000664), CX3CR1-GFP (JAX 005582), B2m^-/-^(JAX 002087), PMEL-1 (JAX 005023) and OT-I (JAX 003831) mice were purchased from The Jackson Laboratory. P14 mice were kindly donated by the Matthew Spitzer laboratory (UCSF).

### GFAP-minigene mice

The minigene cassette (containing the GFAP promoter sequence, Kozak Sequence, tissue plasminogen activator (tPA) signal peptide sequence, the minigene peptide sequences connected with furin cleavage sites and rabbit ß-globin polyadenylation signal) was inserted in reverse orientation into intron 1 of the ROSA26 locus by CRISPR/Cas9. Homology arms flanking the insertion site were PCR-amplified from a BAC and assembled into the targeting vector, which was co-injected with Cas9 protein and a gRNA (Sequence: CTCCAGTCTTTCTAGAAGAT-GGG) into fertilized C57BL/6 embryos. Founders were identified by PCR and confirmed by sequencing; homozygous lines were established.

### EGFRvIII CAR transgenic mice

CAR knock-in (Ki) mice (*46*) were crossed to CD4-Cre (JAX 022071). Offspring were genotyped for both transgenes as described (*46*). Breeders and experimental mice were maintained Cre-hemizygous.

### Cell lines

SB28 and SB28-murineEGFRvIII glioma lines were established previously in our lab (*46*). Each was retrovirally transduced with a minigene encoding minimal epitopes from gp100 (MHC I) and LCMV gp61-80 (MHC II). Transduced cells were selected in cRPMI containing 30 µg/mL blasticidin, cloned by limiting dilution, expanded, and screened by PCR. Cells were maintained in RPMI (Gibco) with 10% heat-inactivated FBS and 1% penicillin–streptomycin (Gibco 15070063). Mycoplasma tests were negative. Single-cell clones were derived by limiting dilution, expanded, and screened by PCR to confirm stable integration of the minigene cassette.

### T-cells and co-cultures

#### Transgenic CD8^+^ T-cells

Spleens from transgenic mice were processed with ACK lysis. CD8^+^ T-cells were purified using MojoSort™ Mouse CD8 T-cell Isolation Kit (BioLegend 480035). Naïve T-cells (CD62L^+^ CD44^low^) were used immediately for adoptive transfer or in vitro assays. Activated T-cells were generated by incubating the isolated cells with CD3/CD28 Dynabeads (Gibco 11453D), and hIL-2 (30 IU/mL; NIH) in cRPMI [RPMI-1640 + 10% FBS, penicillin–streptomycin, HEPES, GlutaMAX, non-essential amino acids, sodium pyruvate, and 0.5 mM 2-mercaptoethanol] for 48 hours. Bead ratio and cell removal was performed according to the manufactures protocol.

#### CFSE labeling and T-cell: CD11b^+^ co-culture

T-cells (1×10^6^) were labeled with 5 µM CFSE (BioLegend 423801; 1 µL of 5 mM stock in 1 mL PBS) for 20 min at 37 °C (dark), washed into cRPMI with hIL-2 (30 IU/mL; PeproTech 200-02), and plated at 1×10^5^ cells/well. CFSE-labeled T-cells were co-cultured 1:1 with CD11b^+^ cells (see below). General positive control: CFSE-labeled T-cells + activation beads; general negative control: CFSE-labeled T-cells without beads. After 72 h, beads were removed and cells collected for flow cytometry.

#### CAR T-cells

CD3^+^ T-cells from EGFRvIII CAR transgenic mice (8–12 weeks) were isolated with MojoSort™ Mouse CD3 T-cell Isolation Kit (BioLegend 480031), then activated for 48 h at 1×10^6^ cells/well (24-well plates) with CD3/CD28 Dynabeads (Gibco 11453D), hIL-2 (30 IU/mL; NIH), and mIL-15 (50 ng/mL; PeproTech 210-15) in cRPMI [RPMI-1640 + 10% FBS, penicillin–streptomycin, HEPES, GlutaMAX, non-essential amino acids, sodium pyruvate, and 0.5 mM 2-mercaptoethanol]. Cells were expanded 7–10 days in IL-2/IL-15 and maintained at 0.5–1×10^6^ cells/mL.

#### Isolation of CD11b^+^ brain cells

Brains (≤500 mg/C Tube) from GFAP-minigene or C57BL/6, mice were dissociated with the Adult Brain Dissociation Kit (Miltenyi 130-107-677) on a gentleMACS™ Octo Dissociator with Heaters (Miltenyi 130-096-427). Debris and myelin were removed; RBCs were lysed per kit instructions. Suspensions were kept at 2–8 °C in PBS/0.5% BSA (PB buffer). CD11b^+^ cells were labeled with CD11b MicroBeads (Miltenyi 130-093-634) and enriched on MS Columns (Miltenyi 130-042-201) in an OctoMACS Separator (Miltenyi 130-042-109). Purity/viability were assessed by flow cytometry. CD11b^+^ cells were cultured on 0.01% poly-L-lysine–coated plates in DMEM + 2 mM L-glutamine, 10% FBS, and 1% penicillin–streptomycin.

### Focused ultrasound–mediated BBB opening

BBB opening was performed in mice using an RK-50 preclinical FUS system (FUS Instruments) equipped with a spherically focused 1.65 MHz transducer and integrated passive cavitation detection. Animals were induced with isoflurane (3–4%) and maintained at 1–2% on a heated stage; scalp hair was removed, and the head was coupled to the water bath via a thin membrane with degassed ultrasound gel. Bracco sulfur-hexafluoride microbubbles (Lumason/SonoVue) were administered retro-orbitally (200 µL bolus) 10 s before sonication. Each target received 25 ms bursts at 1 Hz for 120 s. The initial free-field peak negative pressure was ∼0.35–0.40 MPa and was adjusted in closed-loop using acoustic-emissions feedback to maximize stable cavitation while suppressing inertial cavitation. Feedback regulated pressure range was 0.35-0.6 MPa. BBB opening was confirmed intra-procedurally by sustained subharmonic and ultraharmonic rise without broadband growth. Targets were spaced to avoid focal overlap; cavitation thresholds and final drive levels were recorded per target. Sonication was applied at seven target sites encompassing the tumour injection coordinate (2 mm lateral to bregma, 3 mm ventral) as well as adjacent peritumoural regions positioned 1.5 mm rostral and caudal to this site, ensuring coverage of both the tumour core and surrounding tissue. The same coordinates were applied bilaterally in non–tumour-bearing control mice.

### Intracranial tumour model

1×10^4^ SB28 cells (±EGFRvIII; ±minigene) in 2 µL PBS were stereotactically injected into the right hemisphere of anesthetized C57BL/6 mice (day 0) as described (*47*). Tumour burden was assessed by bioluminescence (Xenogen IVIS Spectrum) 10 min after intraperitoneal administration of D-luciferin (1.5 mg; GoldBio). Mice were randomized by body weight and baseline bioluminescence before treatment.

### Lymphodepletion and adoptive transfer

Lymphodepletion prior adoptive T-cell transfer was performed with intraperitoneal administration of cyclophosphamide (3 mg/20g mouse) plus fludarabine (1 mg per 20 g mouse). Adoptive T-cell transfer was performed via retroorbital infusion of T-cells diluted in 200 µL HBSS.

### Immune-adjuvants

Poly-ICLC (Hiltonol) was generously provided by Dr. Andreas M. Salazar (Oncovir) and administered intramuscularly at a dose of 50 µg. Recombinant human interleukin-2 (teceleukin) was kindly supplied by the NIH and delivered via retro-orbital injection at a dose of 5 ×10^4^ Units in 100 µL PBS. When combined with LIPU+MB treatment, immune adjuvants were administered immediately following sonication.

### Flow cytometry of brain-infiltrating leukocytes

Mice were euthanized and perfused transcardially with 20 mL cold PBS. Brain-infiltrating leukocytes were enriched by Percoll gradient (GE 17089101) (*46*). Cells (0.5–1×10^6^) were stained with Zombie Violet (BioLegend 423114) and Fc-blocked with TruStain FcX™ PLUS anti-mouse CD16/32 (BioLegend 156604), then with fluorophore-conjugated antibodies (manufacturer-recommended dilutions). Fluorescence-minus-one controls guided gating. Panels are listed in **Extended Data Table 1**. Data were acquired on an Attune NxT (Thermo Fisher Scientific) or Cytek Aurora (Cytek Biosciences) and analyzed in FlowJo v10.8.1.

### Bone-marrow chimeras

#### CX3CR1-GFP chimeras

CX3CR1^GFP+^ were lethally irradiated (2×6 Gy, 4 h apart) and reconstituted with 1×10^7^ GFP-negative bone-marrow cells from age-matched C57BL/6 donors. To prevent infections after irradiation, mice received amoxicillin–clavulanate in drinking water at 0.375mg/mL from 1 day before irradiation to 14 days after; solution was refreshed every 2–3 days. Four weeks later, flow cytometry confirmed GFP^+^ -CD45^low^ -CD11b^+^ microglia (host-derived) and GFP^-^-donor-derived CD45^high^ CD11b^+^ infiltrates.

#### GFAP-minigene β2M-KO chimeras

GFAP-minigene recipients were irradiated as above and reconstituted with 1×10^7^ β2M-deficient or wild-type C57BL/6 marrow. Amoxicillin–clavulanate was administered as above. Successfully engraftment was confirmed by selective MHC I expression loss on CD45^high^ CD11b^+^ cells in blood and brain.

### Immunohistochemistry

Mice were euthanized and immediately transcardially perfused with phosphate-buffered saline (PBS). Brains were collected, fixed overnight in 4% paraformaldehyde (PFA) at 4°C, and cryoprotected in 30% sucrose for 24 hours. Samples were embedded in cryoprotective compound (Tissue-Tek, Science Service) and stored at -80°C. Coronal cryosections (10 µm) were cut using a cryostat (Leica), mounted on glass slides (three sections per slide), and stored at −20°C. Sections were post-fixed in 4% PFA for 10 min, washed in PBS, and incubated overnight at 4°C in PBS containing 0.01% Triton X-100 and anti-mouse CD16/32 (1:100) to block nonspecific staining. The following primary antibodies were applied overnight at 4°C: anti-CD8a (Alexa Fluor 594, clone 17A2, BioLegend; 1:100) and anti-GFAP (Alexa Fluor 488, clone GA5, Invitrogen; 1:500). After PBS washes, slides were mounted with Fluoromount-G containing DAPI (Invitrogen, Thermo Fisher Scientific) for nuclear counterstaining. Images were acquired using a BC43 confocal microscope (Oxford Instruments).

### LFA-1 blocking experiment

To assess whether LIPU+MB–induced antigen-specific T-cell homing depends on LFA-1–ICAM-1 interactions, 1 × 10^!^ T-cells were preincubated for 30 min at 4 °C with 10 µg/mL anti–mouse LFA-1α (CD11a) blocking antibody (InVivoMab anti–mouse LFA-1α, clone M17/4; Bio X Cell) or the corresponding isotype control (InVivoMab rat IgG2a, κ isotype control, clone 2A3). Subsequently, the pretreated T-cells were infused via retro-orbital injection into recipient mice immediately before LIPU+MB ± adjuvant treatment, and their CNS homing efficiency was analyzed 24 hours later with flow cytometry.

### Bulk RNA-sequencing of *in vivo* brain samples

To perform RNA-sequencing (RNA-seq) on *in vivo* brain tissue samples, mice were euthanized and perfused transcardially with 10 mL of cold PBS. Whole brains were then harvested and preserved in RNAlater (Invitrogen, AM7024) at 4℃. Tissue pieces (20–30 mg) were excised from the right frontal cortex consistently across all specimens. Following tissue homogenization using a QIAshredder (Qiagen, 79656), total RNA was extracted using the RNeasy Mini Kit (Qiagen, 74106) with the RNase-Free DNase Set (Qiagen, 79254), according to the manufacturer’s protocol. RNA integrity was assessed using an Agilent Bioanalyzer 5400 and confirmed to have a RIN (RNA Integrity Number) value of 8.7 or greater (range: 8.7–9.3). RNA quality assessment, library preparation, and sequencing were performed by Novogene (Sacramento, CA). Briefly, mRNA was purified from total RNA using poly-T oligo-attached magnetic beads. After fragmentation, first-strand cDNA synthesis was performed using random hexamer primers, followed by second-strand cDNA synthesis. Libraries were constructed through end repair, A-tailing, adapter ligation, size selection, amplification, and purification. Quality control was conducted using Qubit and real-time PCR for quantification, and Bioanalyzer for size distribution assessment. Finally, quantified libraries were pooled and sequenced on an Illumina NovaSeq X Plus PE150 platform at Novogene, based on effective library concentration and desired sequencing depth.

### Bulk RNA-sequencing data processing and analysis

Quality checking, trimming, and barcode removal were performed using fastp (v0.20.0) with default parameters. The FASTQ reads were aligned to the mouse reference genome mm10 (GRCm38.p6) using STAR (v2.7.9a), with transcriptome annotation guidance from gencode.vM25.annotation.gtf. Sorting and indexing were performed using samtools (v1.21), and gene-level expression counts were quantified using Subread/FeatureCounts (v2.0.8)(*48*). All subsequent computational analyses were conducted using R (v4.4.2). Differential expression (DE) analyses comparing two groups were performed using the DESeq2 R package (v1.32.0)(*49*). Sex was included as a confounding variable in the design matrix. The output of DESeq2 analysis is provided as Excel File named Raw Data_DESeq.

### Statistics

Analyses were performed in R (v4.4.2) and GraphPad Prism (v10.3.0). Two-group comparisons used Welch’s t test or the Mann–Whitney U test, as appropriate. Multiple comparisons were assessed by one-way ANOVA with Dunnett’s post hoc test. Survival benefit was assessed by log-rank (Mantel–Cox) test. All tests were two-sided, and P < 0.05 was considered statistically significant. Exact n, statistical test, and multiple-testing correction are reported in the results section or in the supplementary tables.

### Data visualization

Figures were generated in R (ggplot2 v3.3.6) or GraphPad Prism v10.3.0. Schematic illustrations were created in BioRender. Final assembly and formatting were done in Adobe Illustrator v28.1.

## Data and materials availability

RNA-seq data is deposited in GEO (Accession number: GSE333455). All other data are available in the manuscript or the Supplementary Materials. Mouse lines and cells are available on reasonable request, subject to institutional MTA approval.

## Author contributions

Conceptualization: MG, HO

Methodology: MG, AY, TN, AS, PC, MC, AMS, HO

Investigation: MG, AY, VAA, TN, LP, JL, SP, HLB,AZ, KO, PBW, JH, SL,

Visualization: MG, HO

Funding acquisition: MG, HO

Project administration: MG, HO

Supervision: HO

Writing – original draft: MG, HO

Writing – review & editing: All authors.

## Funding support

NIH/NINDS: R35NS105068 (HO)

German Research Foundation: GA 3535/1-1 (MG)

Parker Institute for Cancer Immunotherpay (HO, MG)

Focused Ultrasound Foundation: CA-0248769 (MG,HO)

## Acknowledgments

We thank Dr. Matthew Spitzer for the donation of P14 mice. We thank the Laboratory for Cell Analysis of UCSF Hellen Diller Family Comprehensive Cancer Center, The Flow Cytometry Coreof the Gladstone Institute and the UCSF Preclinical Therapeutics Core for excellent technical assistance. The schematic illustrations were generated with biorender.com.

## Extended Data figures

**Extended Data Fig. 1.**
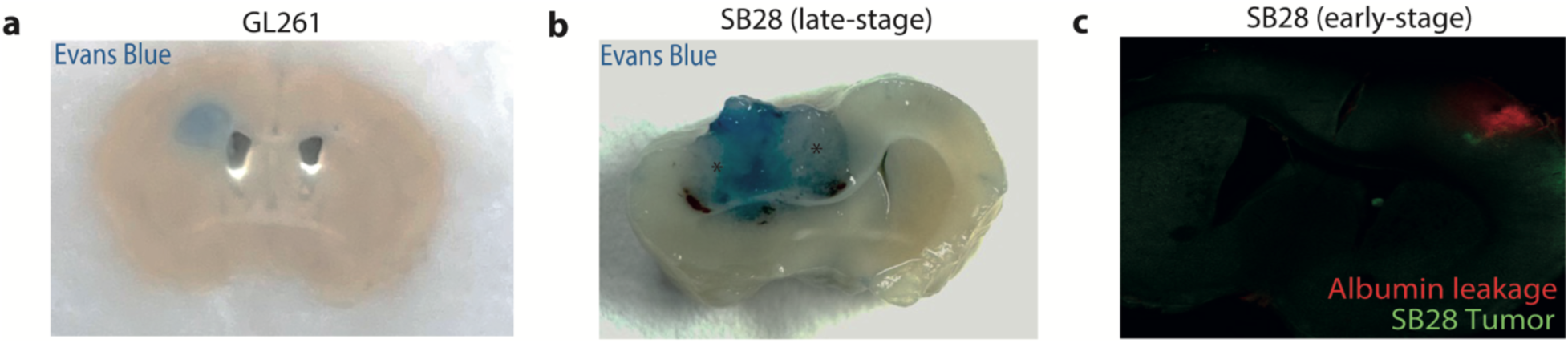
BBB disruption in commonly used glioma models. **a,** Representative coronal brain sections from mice bearing GL261 tumors following intravenous Evans Blue administration. Evans Blue extravasation (blue) reveals heterogeneous blood–brain barrier (BBB) disruption, with pronounced leakage in central tumor regions (dark blue), whereas peripheral tumor areas (*) largely retain BBB integrity. **b,** Representative coronal brain sections from mice bearing SB28 tumors following intravenous Evans Blue administration, showing a similar pattern of heterogeneous BBB disruption with regional variability in vascular permeability. **c,** Immunofluorescence image of SB28 tumors showing focal BBB disruption, visualized by albumin extravasation (red), in proximity to GFP⁺ tumor cells (green). Regions of albumin leakage delineate sites of vascular permeability and local BBB breakdown within the tumor microenvironment.

**Extended Data Fig. 2.**
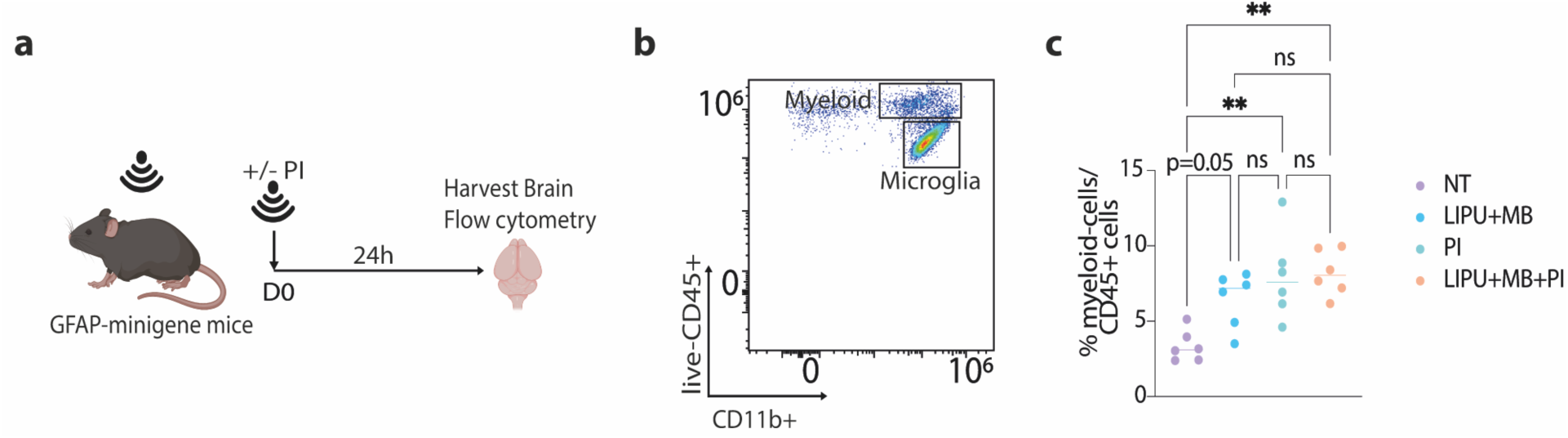
LIPU+MB and poly-ICLC promote recruitment of myeloid cells to the brain. **a,** Experimental design. GFAP–minigene mice received LIPU+MB with or without immune adjuvants (poly-ICLC plus IL-2; PI) on day 0. Brains were harvested 24 h later and analyzed by flow cytometry to assess immune cell infiltration. **b,** Representative flow cytometry gating strategy identifying brain myeloid populations. Live CD45⁺ cells were gated, followed by discrimination of CD11b⁺CD45^high^ infiltrating myeloid cells and CD11b⁺CD45^intermediate^ microglia. **c,** Quantification of myeloid cells among live CD45⁺ brain leukocytes across treatment groups (NT, no treatment; LIPU+MB; PI; LIPU+MB+PI). Each dot represents one mouse. Statistical significance was assessed by one-way ANOVA with multiple comparisons (**P < 0.01; ns, not significant).

**Extended Data Fig. 3.**
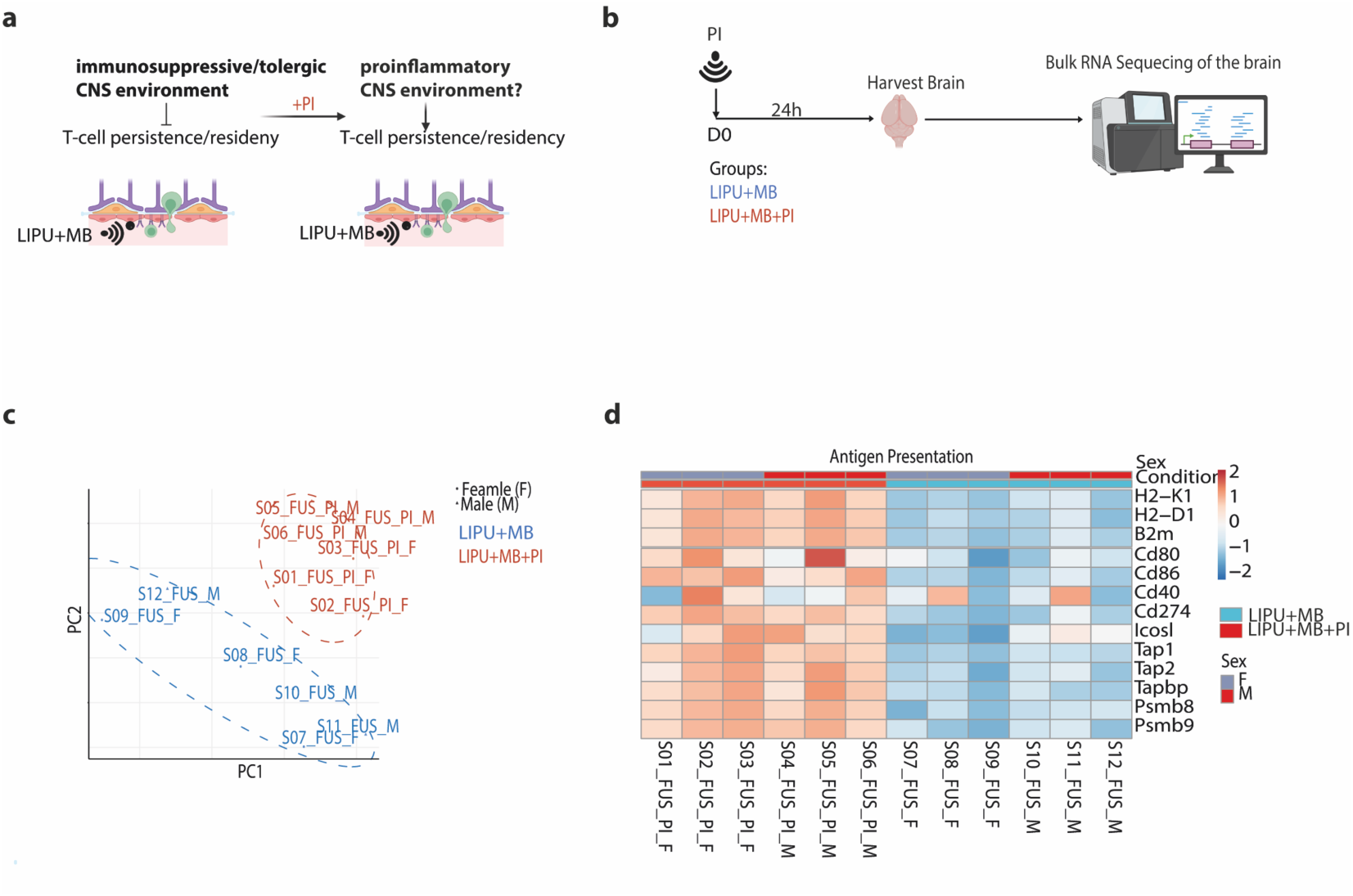
PI promotes proinflammatory transcriptional programs associated with antigen presentation following ultrasound-mediated BBB opening. **a,** Conceptual framework. LIPU+MB-mediated BBB opening occurs within a potentially tolerogenic CNS environment that may limit durable T-cell persistence. Co-administration of immune adjuvants (poly-ICLC plus IL-2; PI) is hypothesized to shift the local milieu towards a proinflammatory state that supports T-cell persistence and residency. **b,** Experimental design. Mice received LIPU+MB with or without PI on day 0. Brains were harvested 24 h later and subjected to bulk RNA sequencing to assess transcriptional responses. **c,** Principal component analysis (PCA) of brain transcriptomes showing separation between treatment groups, indicating distinct transcriptional programs induced by LIPU+MB (blue) and LIPU+MB+PI (red). Each point represents one mouse; dashed ellipses indicate group clustering. **d,** Heat map of differentially expressed genes associated with antigen presentation and immune activation. Brains from mice treated with LIPU+MB+PI show increased expression of genes involved in antigen processing and presentation (for example, H2-K1, H2-D1, Tap1, Tap2 and Psmb8/9) and myeloid activation (for example, Cd80, Cd86 and Cd40) compared with LIPU+MB alone. Annotation bars indicate treatment condition and sex. Colour scale represents row-scaled gene expression (z-score).

**Extended Data Fig. 4.**
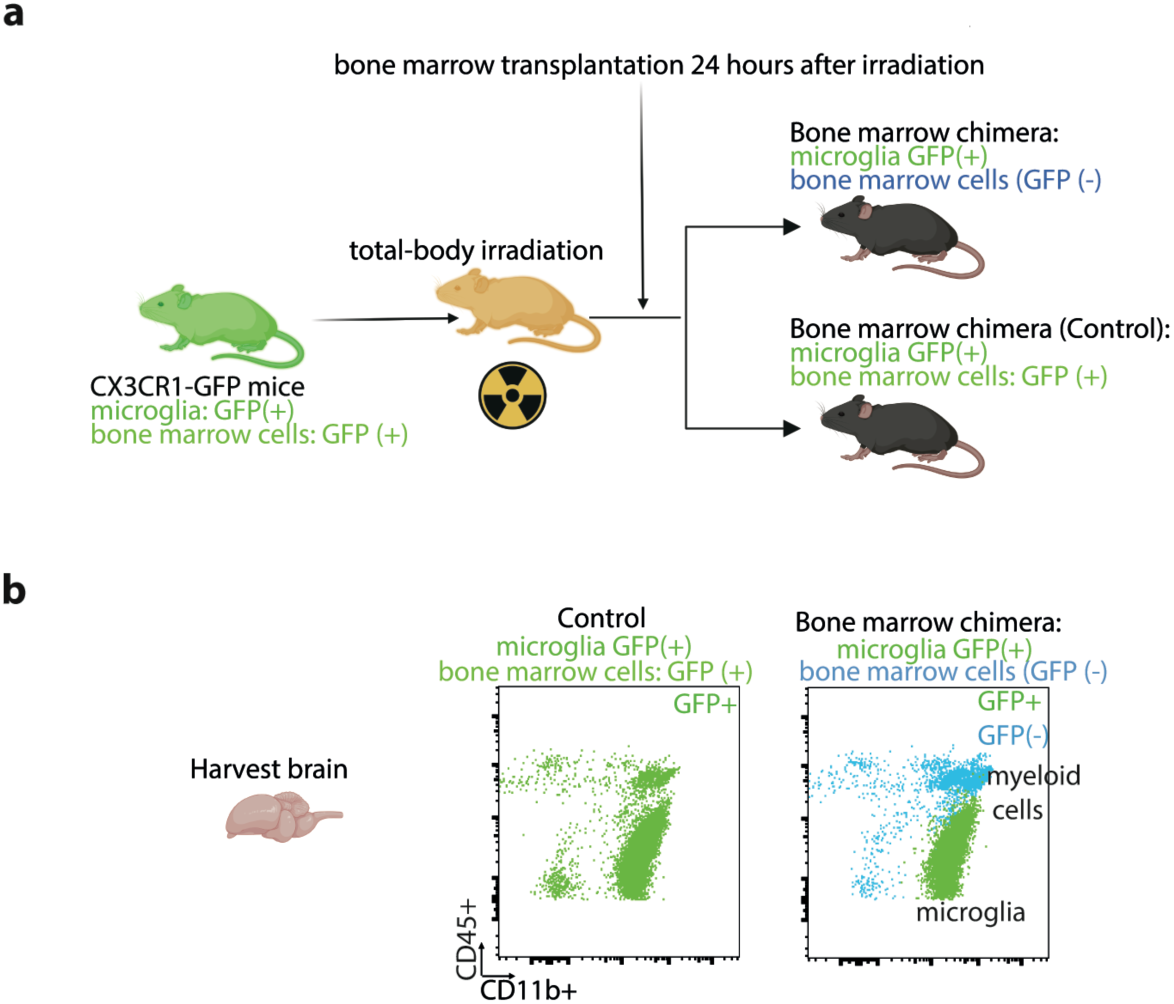
Generation and validation of bone marrow chimeric mice to distinguish CNS-resident microglia from bone marrow–derived cells. **a,** Schematic of the bone marrow chimera approach. CX3CR1–GFP mice, in which CNS-resident microglia and bone marrow–derived cells constitutively express GFP, were lethally irradiated (2 × 5 Gy total-body irradiation, 24 h apart) to ablate endogenous hematopoiesis. Twenty-four hours after the final irradiation, mice received intravenous transplantation of 5 × 10⁶ GFP⁻ bone marrow cells from C57BL/6 donors. As a control, irradiated wild-type mice received 5 × 10⁶ GFP⁺ bone marrow cells from CX3CR1–GFP donors. **b,** Representative flow cytometry plots of brain single-cell suspensions collected 8–10 weeks after transplantation. In mice reconstituted with GFP⁻ bone marrow, CNS-resident microglia retained GFP expression (GFP⁺), whereas infiltrating bone marrow–derived cells were GFP⁻, enabling discrimination between these populations. In control chimeras, bone marrow–derived cells were GFP⁺, confirming efficient engraftment. This system enables fate mapping of radio-resistant resident microglia and radio-sensitive bone marrow–derived myeloid cells within the CNS.

**Extended Data Fig. 5.**
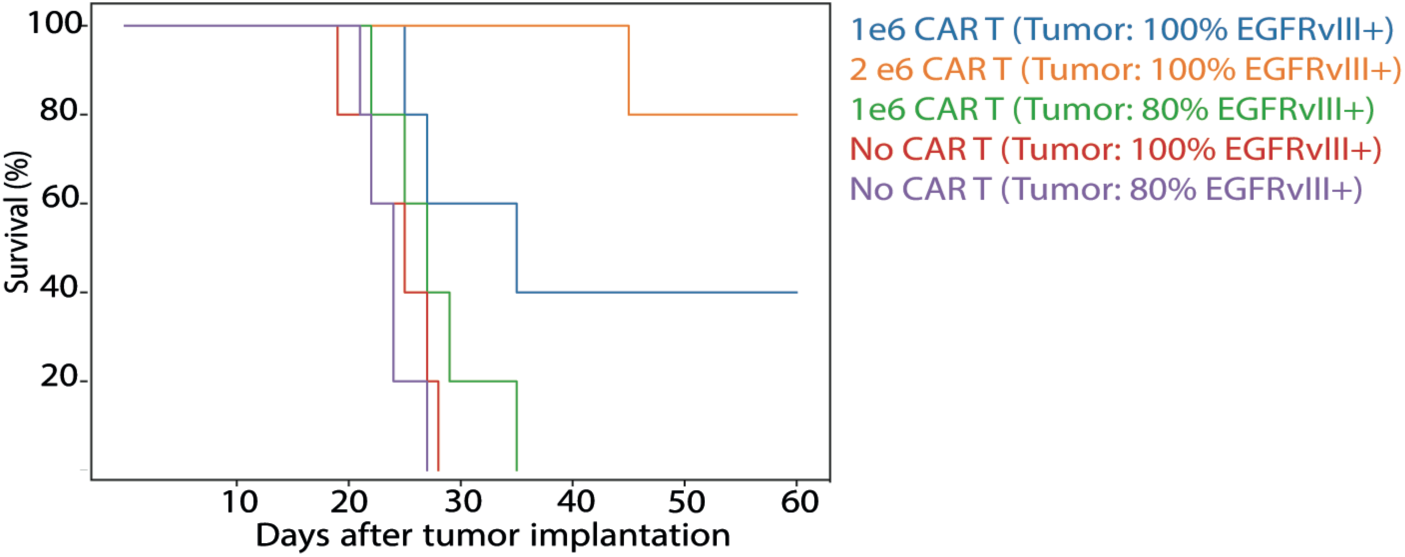
Efficacy of EGFRvIII-directed CAR T cells in homogeneous versus heterogeneous SB28 gliomas. Kaplan–Meier survival analysis showing therapeutic efficacy of EGFRvIII-directed CAR T cells. In mice bearing homogeneous tumors (100% EGFRvIII⁺), CAR T-cell therapy prolonged survival in a dose-dependent manner, with 2 × 10⁶ cells providing the greatest benefit. In contrast, CAR T-cell treatment did not extend survival in mice bearing heterogeneous tumors (80% EGFRvIII⁺), consistent with antigen escape–mediated resistance. Control mice (PBS) exhibited rapid tumor progression irrespective of EGFRvIII expression.

**Extended Data Fig. 6.**
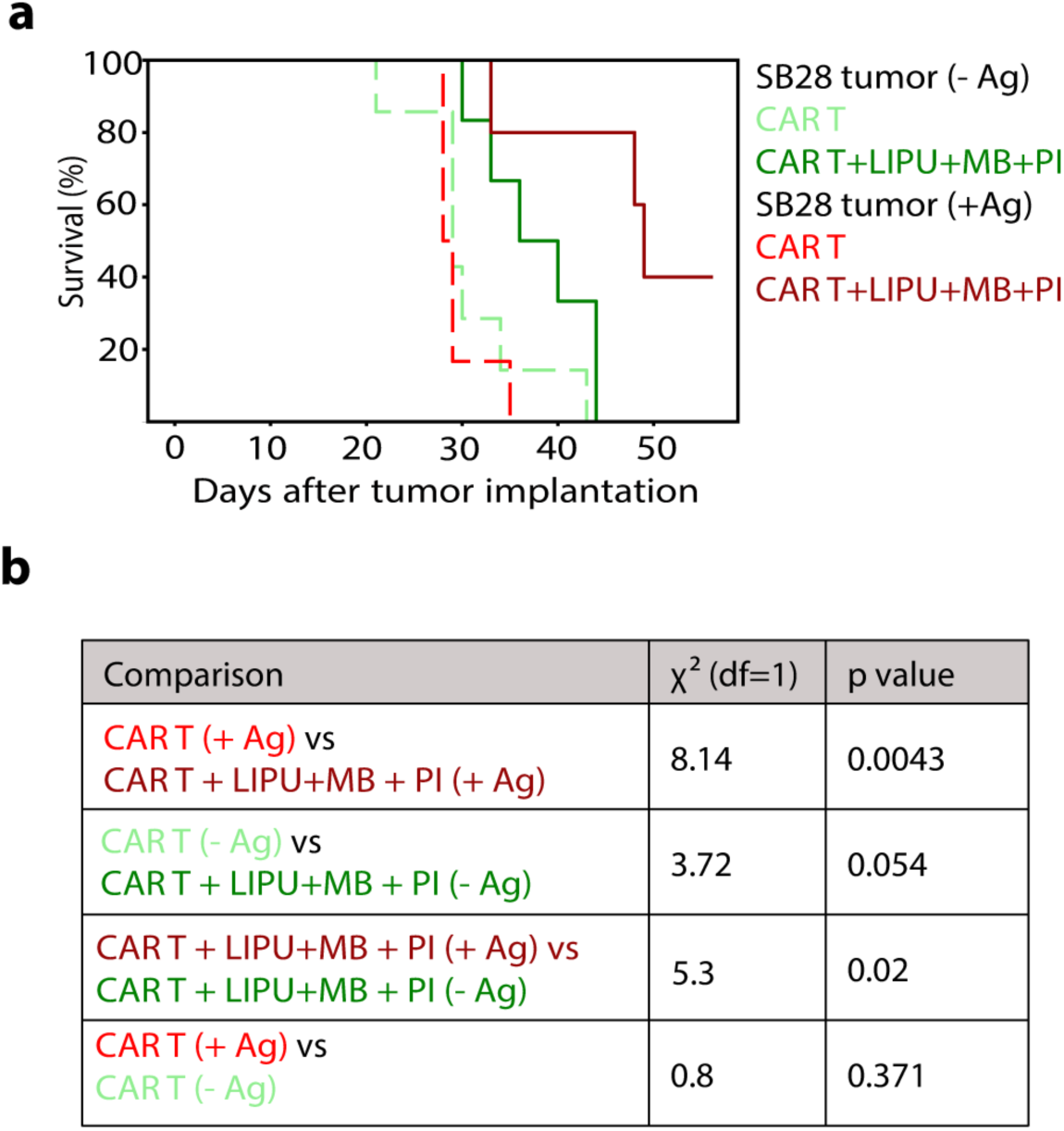
Antigen-dependent efficacy of combined BBB modulation and immune activation. **a,** Kaplan–Meier survival analysis of mice bearing SB28 tumors expressing gp100 and LCMVgp61–80 antigens (+Ag) or lacking antigen expression (−Ag), following treatment with CAR T cells alone or in combination with LIPU+MB+PI. **b,** Statistical analysis of survival differences using the log-rank (Mantel–Cox) test.

## Supplementary tables

**Extended data table 1.**
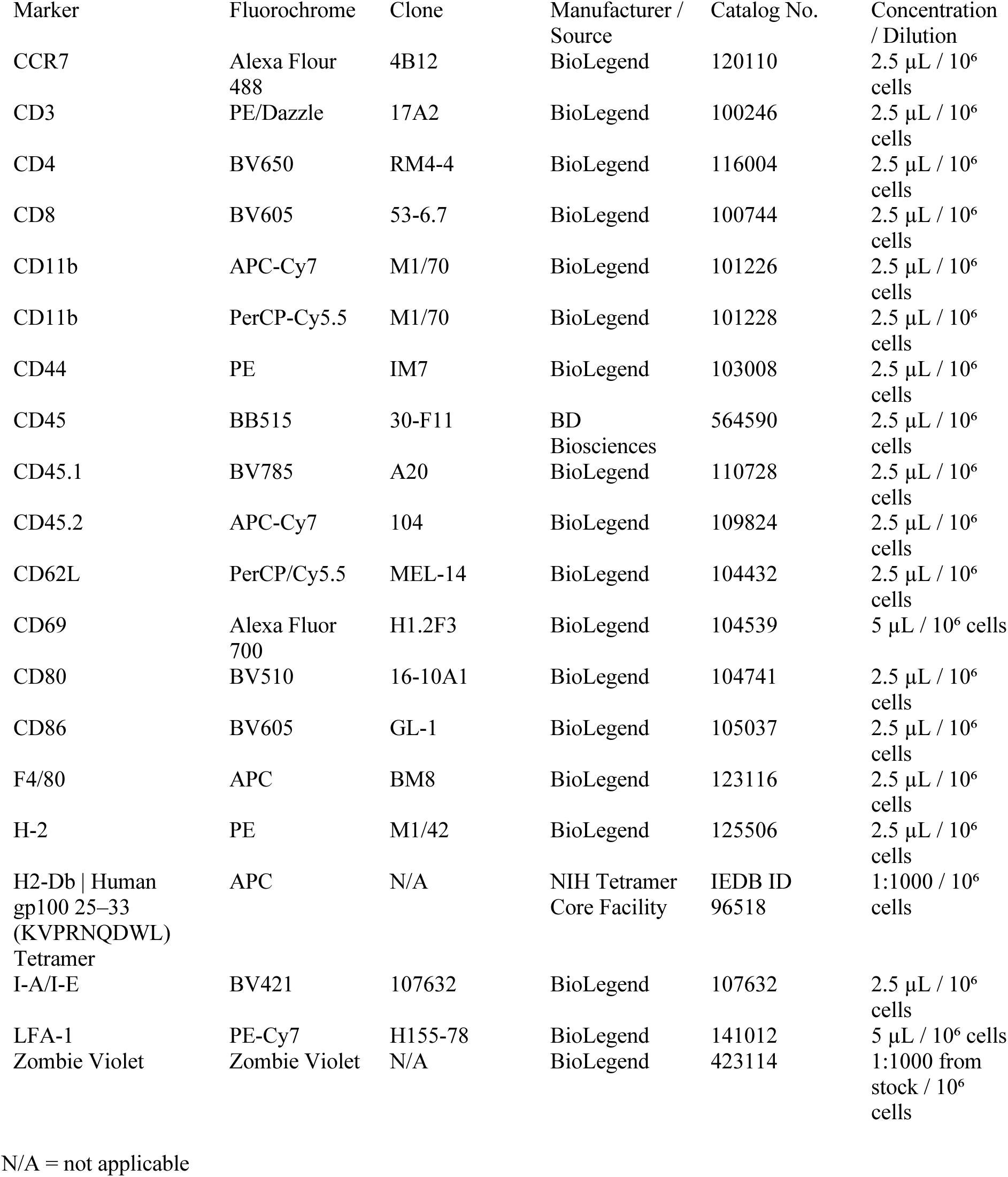
Antibodies and Tetramers Used in Flow Cytometry Analyses.

